# Modulation of TRPV4 Protects against Degeneration Induced by Sustained Loading and Promotes Matrix Synthesis in the Intervertebral Disc

**DOI:** 10.1101/2022.11.18.516994

**Authors:** Garrett W.D. Easson, Alireza Savadipour, Akila Anandarajah, Leanne E. Iannucci, Spencer P. Lake, Farshid Guilak, Simon Y. Tang

**Author notes:** **Corresponding author**, Simon Tang, Ph.D., MSCI, 660 S Euclid Ave, Campus Box 8233, St. Louis, MO 63110.

## Abstract

While it is well-known that mechanical signals can either promote or disrupt intervertebral disc (IVD) homeostasis, the molecular mechanisms for transducing mechanical stimuli are not fully understood. The transient receptor potential vanilloid 4 (TRPV4) ion channel activated in isolated IVD cells initiates extracellular matrix (ECM) gene expression, while TRPV4 ablation reduces cytokine production in response to circumferential stretching. However, the role of TRPV4 on ECM maintenance during tissue-level mechanical loading remains unknown. Using an organ culture model, we modulated TRPV4 function over both short-(hours) and long-term (days) and evaluated IVDs’ response. Activating TRPV4 with the agonist GSK101 resulted in a Ca^2+^ flux propagating across the cells within the IVD. NF-κB signaling in the IVD peaked at 6 hours following TRPV4 activation that subsequently resulted in higher IL-6 production at 7 days. These cellular responses were concomitant with the accumulation of glycosaminoglycans and increased hydration in the nucleus pulposus that culminated in higher stiffness of the IVD. Sustained compressive loading of the IVD resulted in elevated NF-κB activity, IL-6 and VEGF-A production, and degenerative changes to the ECM. TRPV4 inhibition using GSK205 during loading mitigated the changes in inflammatory cytokines, protected against IVD degeneration, and but could not prevent ECM disorganization due to mechanical damage in the annulus fibrosus. These results indicate TRPV4 plays an important role in both short-and long-term adaptations of the IVD to mechanical loading. The modulation of TRPV4 may be a possible therapeutic for preventing load-induced IVD degeneration.

## Introduction

Low back pain (LBP) is one of the most prevalent musculoskeletal diseases in the U.S., and it is the leading cause of disability.^1–3^ The degeneration of the intervertebral disc (IVD), the soft cartilaginous tissue between the vertebrae, is a significant contributor to low back pain.^4–6^ The IVD experiences continuous mechanical loading and unloading cycles that enables motion and load transmission in the skeleton. While mechanical stimulation is crucial for IVD homeostasis and maintenance, over-loading leads to tissue damage with the subsequent expression of inflammatory cytokines such as interleukin 1β (IL-1β), IL-6, and IL-8 in IVD cells.^7–12^ Although the transient expression of inflammatory cytokines such as IL-6 have an essential regulatory role in tissue regeneration, fracture repair, and wound healing,^13–15^ their sustained expression can eventually cause the release of catabolic enzymes, initiate immune cell recruitment, and provoke vascular and neurite invasion of the IVD.^8,16–19^

Mechanosensitive ion channels are activated by mechanical stimuli and allow for the transport of ions across the cell membrane, and are hypothesized to participate in the IVD’s responsiveness to its loading environment.^20,21^ One candidate mechanosensitive channel in the IVD is transient receptor potential vanilloid 4 (TRPV4), a nonselective cation channel in the TRP family of channels.^22,23^ TRPV4 is an important osmo-and mechano-regulator in a variety of cell types, including chondrocytes, osteocytes, cardiovascular endothelial cells, and renal epithelial cells.^24–26^ TRPV4 is crucial for mechanosensing in articular cartilage and has been shown to regulate the anabolic response of chondrocytes to mechanical loading.^27–29^ Similar to articular cartilage in both its motion-enabling joint function and matrix constituents, the mechanosensing mechanisms may be conserved in the IVD. Moreover, the application ofload to both cartilage and the IVD causes perturbations in the local osmotic environment that triggers a Ca2_+_ signaling response.^30 31^ However, it is unknown whether this response is mediated by TRPV4.

In IVD cell culture, inhibiting TRPV4 reduces the hypo-osmotic-mediated production of IL-1β and IL-6, and the production of IL-6 and IL-8 mediated by high-magnitude strains.^32,33^ Conversely, activation of TRPV4 in IVD cells promotes the expression of matrix protein genes including ACAN and COL2A1.^34^ We therefore hypothesized that TRPV4 regulates the mechanotransduction of physiologically relevant loading in the IVD. To investigate the complex interactions between physiologic loading, maintenance of the extracellular matrix (ECM), and modulation of TRPV4, we used an organ culture IVD system that allowed us to apply compression in a sustained manner, in the presence or absence of a TRPV4 agonist or antagonist, and observe outcomes on the entire IVD.

We first examined the effects of transient TRPV4 activation using a small molecule agonist (GSK1016790A, i.e., GSK101) on the cellular level ca across the IVD. We then compared the IVDs’ adaptations resulting from prolonged activation of TRPV4 and corresponding hypo-osmotic conditions. Finally, we evaluated the role of TRPV4 under sustained compression of the whole IVD at physiologically relevant loads. Our results show that the transient, single-bout activation of TRPV4 in IVD organ culture increases the intracellular Ca^2+^ flux and a cascade of NF-κB signaling and production of IL-6. The repeated activation of TRPV4 over a one-week period improved the compressive stiffness of the IVD and increased glycosaminoglycan GAG content and hydration in the nucleus pulposus (NP). Further, the inhibition of TRPV4 during the repetitive hypo-osmotic conditions blunted IL-6 production by the IVD. The sustained mechanical compression of IVDs resulted in degeneration along with prolonged expression of IL-6 and vascular endothelial growth factor A (VEGFA). The inhibition of TRPV4 during this sustained loading alleviated the loading-induced degeneration and protected from the increases of IL-6 and VEGFA, but it did not prevent the load-induced collagen disorganization in the annulus fìbrosus. Taken together, the transient activation of TRPV4 initiates the cascade of Ca^2+^ signaling, NF-κB signaling, and IL-6 expression, with the repeated activation of TRPV4 increasing GAG accumulation and the stiffness of the IVD ECM. Inhibition of TRPV4 mitigated the degenerative effects of sustained mechanical loading. These results suggest that modulation of TRPV4 could provide a novel approach for preventing degeneration of the IVD following deleterious loading.

## Materials and Methods

### 2.1 Animals

All animal experiments were performed in compliance with the Washington University in St. Louis Institutional Animal Care and Use Committee. Sixteen-week old Mice carrying the NF-κB-luciferase reporter gene (FVB.Cg-Tg(HIV-EGFP,luc)8Tb/J; JAX) were used. The mice were euthanized with CO_2_ for two minutes and then disinfected in 70% ethanol. The whole lumbar spine was then removed and separated into three functional spine units (FSUs): Ll/L2, L3/L4, L5/L6. All excess tissue including the spinal cord was removed. FSUs were immediately placed in culture media in 24-well plates and pre-conditioned for seven days prior to all experiments. Culture media consisted of 1:1 Dulbecco’s modified Eagle’s medium: Nutrient mixture F-12 (DMEM:FI2) supplemented with 20% fetal bovine serum and 1% penicillin-streptomycin.^35^ The FSUs were cultured under the following conditions (37°C, 5% CO_2_, 100% humidity) and were assigned to one of three distinct experiments: transient activation of TRPV4, repeated activation of TRPV4, and sustained activation of TRPV4 (Figure 1). Our prior work demonstrated that these culture conditions maintains the viability of all cells in the IVD, including those in the cartilaginous end-plate, but not cells in the bone marrow space or osteocytes.^35^

**Figure 1:**
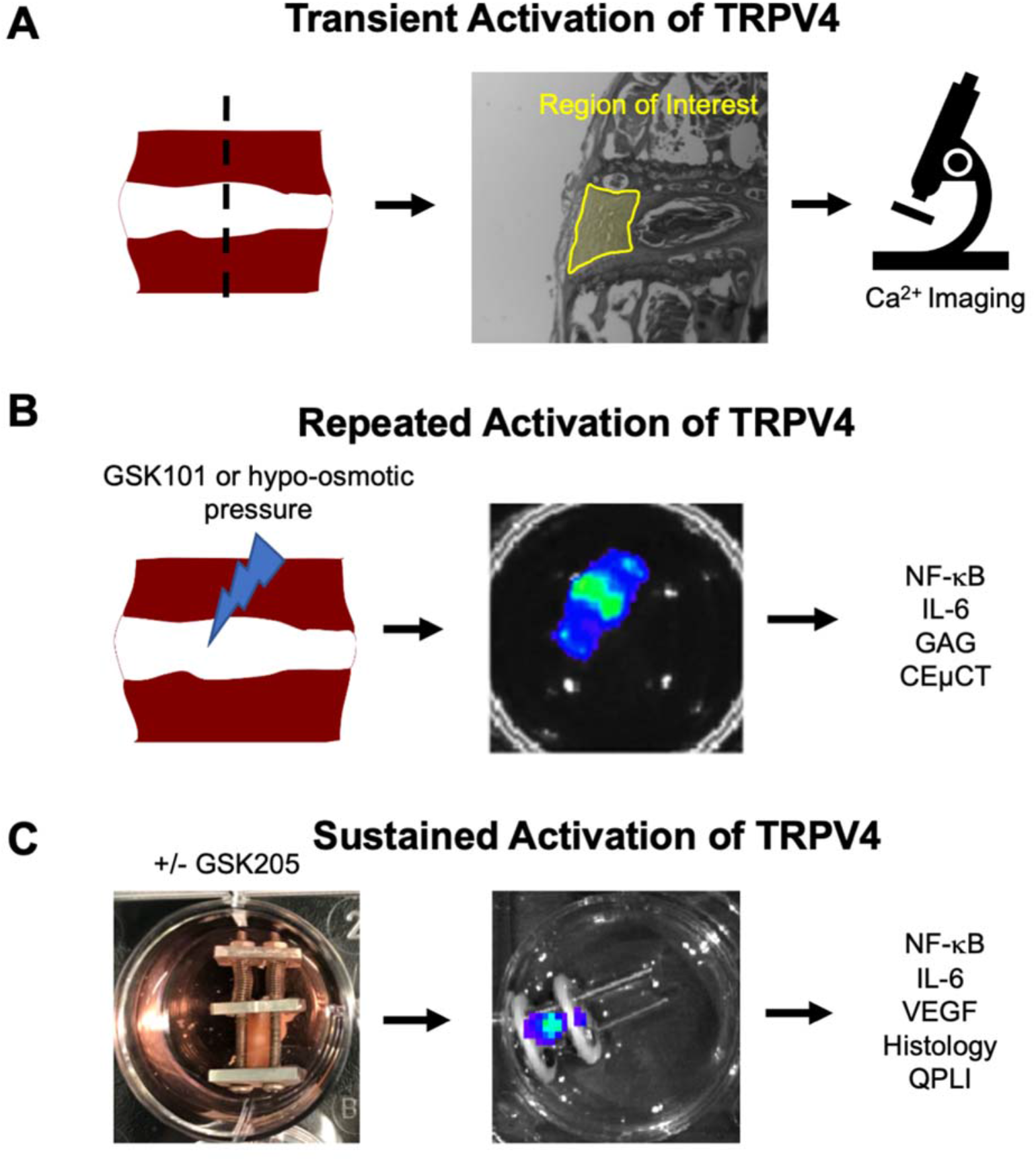
Functional Spine Units including the IVD were extracted from sixteen-week old mice carrying the NF-κB-luciferase reporter transgene, and then were divided into 3 sets of organ culture experiments: (A) Observation of calcium flux following a single, transient bout of TRPV4 activation. Fluorescent Ca^2+^ signals were observed using confocal microscopy immediately following GSK101 administration (n=3); (B) the repeated activation of TRPV4 using GSK101 (n=15) or by hypo-osmotic pressure (n=8); and (C) the sustained activation through static compression throughout the culture period (n=11). The outcome measures utilized in (B) and (C) include NF-κB-luciferase activity, IL-6 and VEGF cytokine production measured by ELISA, glycoaminoglycan (GAG) composition measured by the dimethyl-methylene blue (DMMB) assay, IVD structure measured by contrast-enhanced μCT (CEμCT), IVD degeneration measured by histologic grading, and IVD tissue organization measured by quantitative polarized light imaging (QLPI).

### 2.2 Confirmation of TRPV4 function through Ca^2+^ signaling

TRPV4 is a Ca^2+^ preferential ion channel, and its activation results in fluxes of intracellular Ca^2+^. To confirm that TRPV4 is present and functional in our functional spine unit (FSU) cultures (n=3), we measured changes in intracellular Ca^2+^ concentrations using ratio imaging of Fluo-4 and Fura Red Ca^2+^ indicators. The FSUs were incubated with Fluo-4 and Fura Red Ca^2+^ indicators for 1 hour. Changes in intracellular Ca^2+^ were observed in real-time (Zeiss LSM 880) and the ratio of Fura Red intensity (decreases with low Ca^2+^) to Fluo-4 intensity (increases with high Ca^2+^) were computed. The Ca^2+^ gradient was quantified using a custom MATLAB script by dividing the green channel intensity by the red channel intensity frame-by-frame to obtain relative fluorescent units (RFU). The fluorescence was observed using confocal microscopy at ~150μm of depth into the annulus fibrosus (AF) to evaluate the activation of TRPV4 in AF cells in their native extracellular matrix. DMSO was added to the IVD at the start of the live imaging session, and then cells were monitored for approximately 200s. Following this period, 1μM GSK101 was added to the culture to agonize TRPV4. Concentrations of 100μM and ImM of GSK101 were subsequently added to the culture media of respective IVDs with each IVD receiving just one concentration to determine the dose-dependent response.

### 2.3 Repeated TRPV4 activation using the GSK101 agonist in FSU culture

FSUs in culture were either incubated with DMSO (n=12) or the TRPV4 agonist GSK101 (2μM; n=14) for seven consecutive days for three hours each day. Continuous high intracellular Ca^2+^ concentrations are toxic for cells, and sustained Ca^2+^ saturation can induce apoptosis. Thus, we selected a dose of GSK101 that activates, but does not saturate intracellular Ca^2+^ signaling.^36^ All treatment was delivered in ImL of fresh culture media containing either DMSO or GSK101. After the three hours, the IVDs were returned to fresh media under the prescribed conditions.

### 2.4 Real-time observation of NF-κB signaling

NF-κB signaling was observed longitudinally for several experiments. Luciferin (30mg/mL) was added to the culture media, which then binds to the luciferase transgene to produce detectable bioluminescence quantifying the degree of NF-κB signaling in the IVD.

Bioluminescence was measured 30 minutes following luciferin addition at either: Time 0 hr, 3 hr, 6 hr, 9 hr, or 24 hr for the short-term study; or Days 0, 3, 7, and 14 of the organ culture for the longer term study. Samples were imaged using an IVIS Imaging System (Xenogen Corp.) at a 10s exposure time. Lipopolysaccharide (LPS) was used to produce a positive NF-κB control (1μg/mL, Sigma Aldrich: L8274). The Day 0 time point is the baseline prior to the application of experimental conditions. Following every imaging timepoint, the FSUs were transferred to fresh media.

### 2.5 Cytokine production by measured ELISA

Media was collected from the FSU cultures on Days 0, 3, 7, and 14 of the primary study and stored at −80°C. The concentrations of IL-6 and VEGFA were determined using an enzyme-linked immunosorbent assay (Mouse IL-6 ELISA kit, ThermoFisher; Mouse VEGF-A DuoSet ELISA kit, R&D Systems). All media was collected two days following media change, such that measured cytokine concentration reflected two days of production.

### 2.6 Contrast-enhanced micro-computed tomography

To evaluate structural changes, IVDs were imaged using contrast-enhanced micro-computed tomography (CEμCT) with ioversol as the contrast agent.^37^ Intact IVDs were incubated in 50% Ioversol (OptiRay 350, Guerbet Pharma) at 37°C for 8 hours prior to imaging. Following incubation, samples were scanned using Scanco40 microCT system (Zurich, Switzerland) at 45 keVp, 177 μA, 10.5μm voxel size, and 300 ms integration.

### 2.7 Mechanical behavior of the whole IVD

Mechanical behavior of the IVDs was quantified using displacement-control dynamic compression testing (BioDent, Active Life Scientific).^38^ After FSUs were adhered to aluminum platens, disc height was determined by calipers and used to determine the input strain values. Samples were placed in a phosphate buffered saline bath and preloaded to 0.02N. A sinusoidal compressive waveform was applied at 5% strain at 1Hz for 20 cycles. Average stiffness was determined from the second through the final loading cycles.

### 2.8 GAG quantification by DMMB assay

Dimethylmethylene blue (DMMB) assay was used following the culture period to measure the total GAG content in a subset of tested IVDs. IVDs were isolated, massed, and then digested in papain overnight in 65°C. Supernatant of samples were collected following centrifuging and plated alongside chondroitin sulfate standards (Sigma Aldrich). 250μL of DMMB binding dye was added to each sample and samples were read at an absorbance of 525nm using a spectrophotometer.

### 2.9 Repeated hypo-osmolar exposure

FSUs underwent a 7-day preconditioning period in 400 mOsm media (representing approximately standard culture conditions) following extraction. FSUs were then placed in either a 400 mOsm or 200 mOsm (hypo-osmolar) media, with either DMSO (vehicle) or 10μM GSK205, for 3 hours each day for 5 consecutive days. Osmolarity was titrated by adding sucrose to culture media.

### 2.10 Sustained static loading of the functional spine units in culture

A custom platen-spring device was developed for loading the FSU and evaluating the role of TRPV4 in the load-mediated response of the IVD. The device utilized machined platens coupled with springs of known spring constants to deliver controlled loads based on the distance between the two platens. FSUs were statically loaded for 24 hours at ~O.2MPa, which is approximately two times the body weight of a mouse (~25g) applied across the area of a lumbar IVD.^39^ DMSO (vehicle) or GSK205 (TRPV4 antagonist) were added to the medium of compressed IVDs during the entire duration of loading. Lipopolysaccharide (LPS) was added to a subset of IVDs for the positive controls for inflammation. Following the 24 hours of loading, the device was removed and FSUs were cultured for 14 days. Following the culture period, samples were processed for histology and media collected at Days 7 and 14 was measured for cytokine production.

### 2.11 Histology and mouse IVD degeneration scoring

All samples were fixed in formalin and processed for paraffin embedding. Samples were then sectioned to 10μm thickness and underwent Safranin O staining with a Fast Green counter stain. IVDs degeneration was quantified using a standardized scoring system.^40^ IVDs were graded by a single grader in a blinded fashion on three non-consecutive days. An intraclass correlation (ICC) was calculated between the scoring days to ensure consistent grading (Supplemental Figure S1).

### 2.12 Quantitative polarized light microscopy

Transmission-mode quantitative polarized light imaging (QPLI) was performed on Safranin-O stained sections of IVD as previously described.^41,42^ In brief, a SugarCube White LED Illuminator (Edmund Optics, Barrington, NJ, USA) with a Dolan-Jenner fiber optic backlight (Edmund Optics, Barrington, NJ, USA) was used to illuminate the histological section of interest. The light passed through a circular polarizing film (Edmund Optics, Barrington, NJ, USA) before being transmitted through the slide. The illuminated section was imaged using a division of focal plane polarization camera (FLIR, Wilsonville, OR, USA) with an achromatic 10x objective. The degree of linear polarization (DoLP) and the angle of polarization (AoP) were calculated on a pixelwise basis for the entire field of view. DoLP corresponds to the strength of collagen fiber alignment and AoP corresponds to the orientation of collagen fibers. Images were manually segmented to select a region of interest spanning the AF. DoLP and AoP color maps were generated within the specified region of interest (ROI) and overlayed on the grayscale image for qualitative analysis. Average (AVG) DoLP and Standard deviation (STD) of DoLP values were calculated within the ROI.

### 2.13 Statistical Analysis

All statistical analysis was performed in either MATLAB or Graphpad Prism 9 software. Comparisons across experimental factors were conducted using student’s t-test or multi-way ANOVA as appropriate, and Tukey’s adjusted method was used for post hoc comparisons between groups. Results were considered statistically significant when p < 0.05.

## Results

### 3.1 TRPV4 activation induces Ca^2+^ signaling in IVD cells

To evaluate the role of TRPV4’s activation in Ca^2+^ signaling, the annulus fibrosis (AF) and nucleus pulposus (NP) cells in the intact IVD were observed with confocal microscopy using ratio of intensities of fluorescent Ca^2+^ indicators. The addition of DMSO had no observable changes in fluorescence intensity (data not shown). The TRPV4 agonist GSK101 was added to the culture media, and then the fluorescent Ca^2+^ signal was first observed in the cells of the outer AF. The fluorescence then propagated radially inward towards the nucleus pulposus (Figure 2A-C; Supplemental Video). The percentage of responsive cells increased in a dose-dependent manner. 1μM of GSK101 elicited the slowest response that activated only 30% of cells (Figure 2D). More than 95% of observed cells experienced Ca^2+^ influx within 600s of adding 100μM GSK101 (Figure 2D). The 1 mM of GSK101 achieved greater than 95% activation in under 120s. As GSK101 is a specific chemical-activator of TRPV4, the observed intracellular Ca^2+^ flux confirms the presence and function of TRPV4 in both AF and NP cells. These Ca^2+^ fluxes typically resolved within an hour of TRPV4 agonist removal.

**Figure 2:**
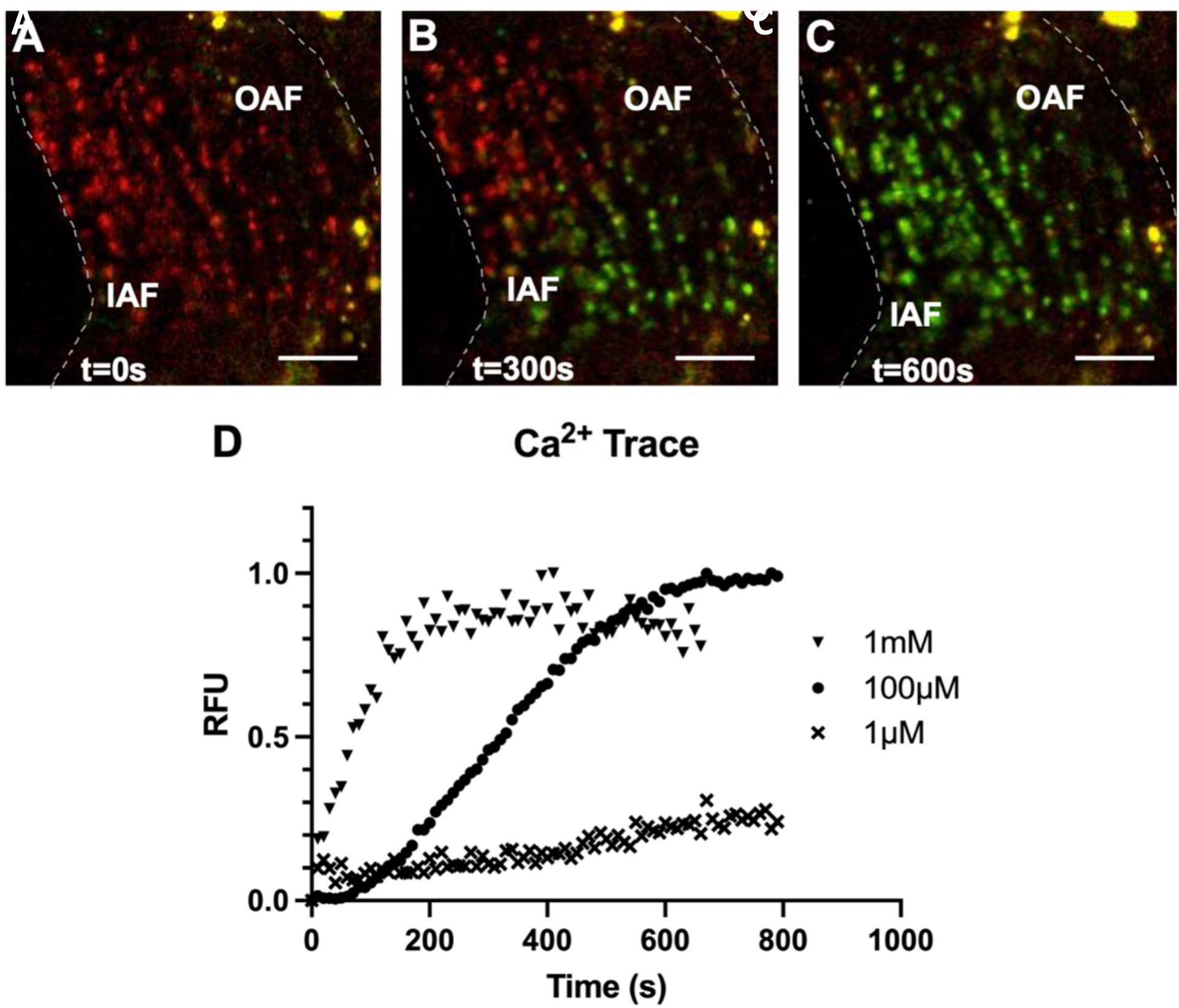
Representative confocal microscopy images showing that activation of TRPV4 by exposing the IVD to GSK101 rapidly increases intracellular Ca^2+^ ions. Images are from the sample treated with 100μM of GSK101. The dotted lines indicate the borders of the annulus fibrosus, with IAF indicating the border of the inner annulus fibrosus, and OAF indicating the border of the outer annulus fibrosus. (A) The red signal indicates low intracellular Ca^2+^. (B) As the GSK101 gradually perfuses into the tissue to open TRPV4, the influx of intracellular calcium across the IVD cells is confirmed by the green fluorescence of the Fluo-4 Ca2+ indicator. (C) Within 600s, the majority of cells show prevalent TRPV4-activation. (D) The time-dependence and dose-dependence of the resultant calcium flux are determined by the ratio between green and red fluorescence intensities at three different GSK101 concentrations. Scale bars are 100μm.

### 3.2 NF-κB signaling is downstream of TRPV4 activation

To evaluate the extent of NF-κB expression, bioluminescence activity of the IVD was measured following a single bout of 3-hour TRPV4 activation by GSK101. The monitoring of the subsequent 24-hour time course revealed a transient peak at 6 hours (p<0.05) that gradually returned to baseline levels by 24 hours post-GSK101 exposure (Figure 3A, B). The timing of the NF-κB peak signal typically lagged the peaks of the Ca^2+^ time-course by an order of magnitude.

**Figure 3:**
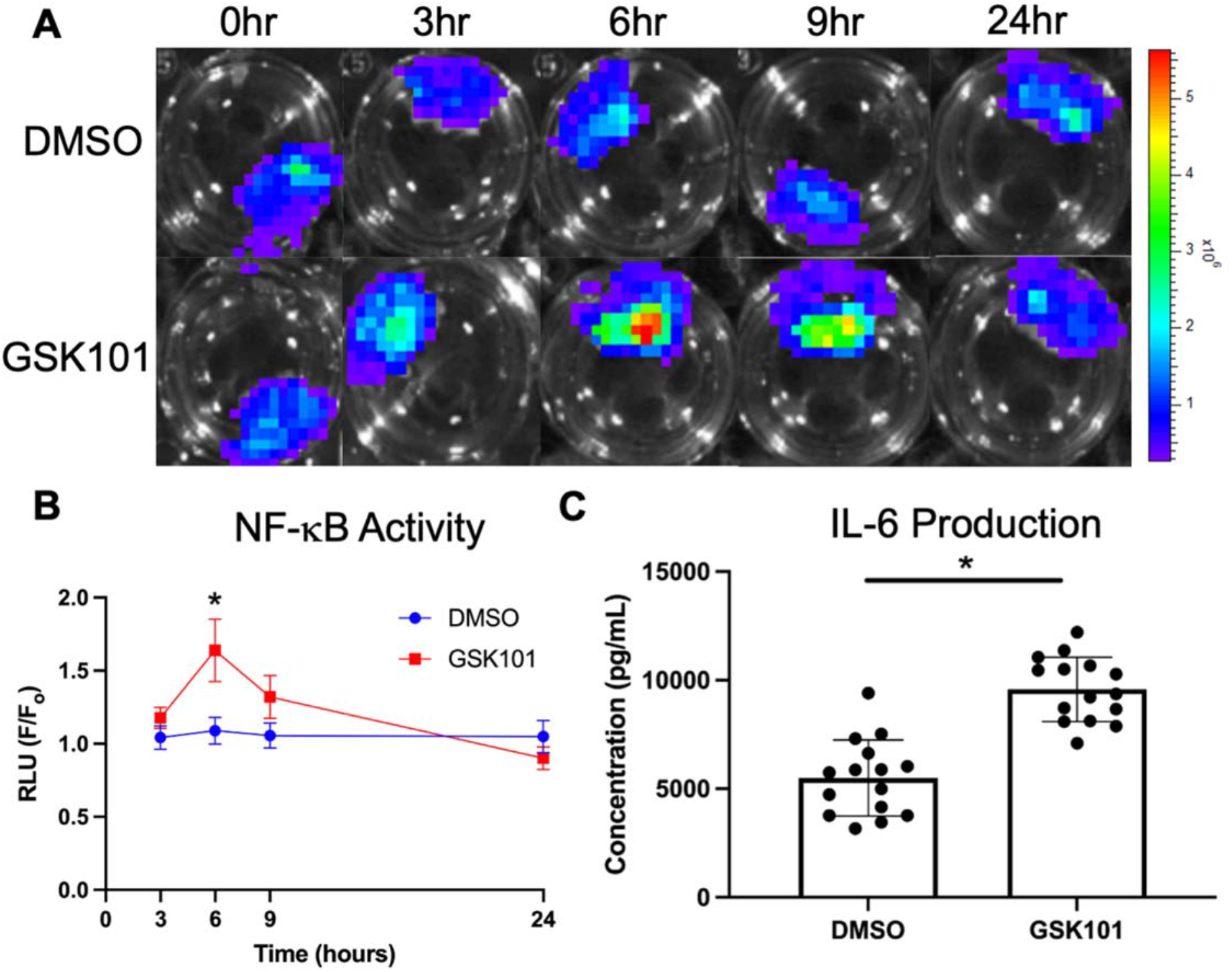
(A,B) NF-κB activity, reported by bioluminescence, increased significantly following GSK101 administration (n=11) compared to DMSO (n=10) at t = 6hr (p<0.05), but this increase completely subsided by 24 hours. (C) In a separate culture experiment (n=15), administration of GSK101 to activate TRPV4 for 3 hours each day for 7 days increased IL-6 production by the IVDs (p<0.05). Data in (B) were statistically analyzed via one-way ANOVA with post hoc Tukey’s HSD where * indicates p < 0.05. Data in (C) were statistically analyzed via t-test where * indicates p< 0.05.

### 3.3 Repeated TRPV4 activation elicits an inflammatory response and augments the ECM

Whereas a singular dose of GSK101 initiates a transient inflammatory response that is resolved rapidly, the repetitive activation of TRPV4 was applied to determine the effects of cytokine production. Seven days of repeated TRPV4 activation increased IL-6 production by the IVDs (Figure 3C) but not VEGFA (p=0.41). Mechanical testing of these IVDs revealed increased compressive stiffness due to TRPV4 activation (Figure 4A). Those IVDs subjected to repeated TRPV4 activation exhibited significantly higher GAG content than the IVDs exposed to DMSO (Figure 4B), demonstrating that this repetitive TRPV4 activation promotes matrix deposition. CEμCT confirmed that the increase in GAG is primarily localized in the NP (Figure 4C, D).^43^

**Figure 4:**
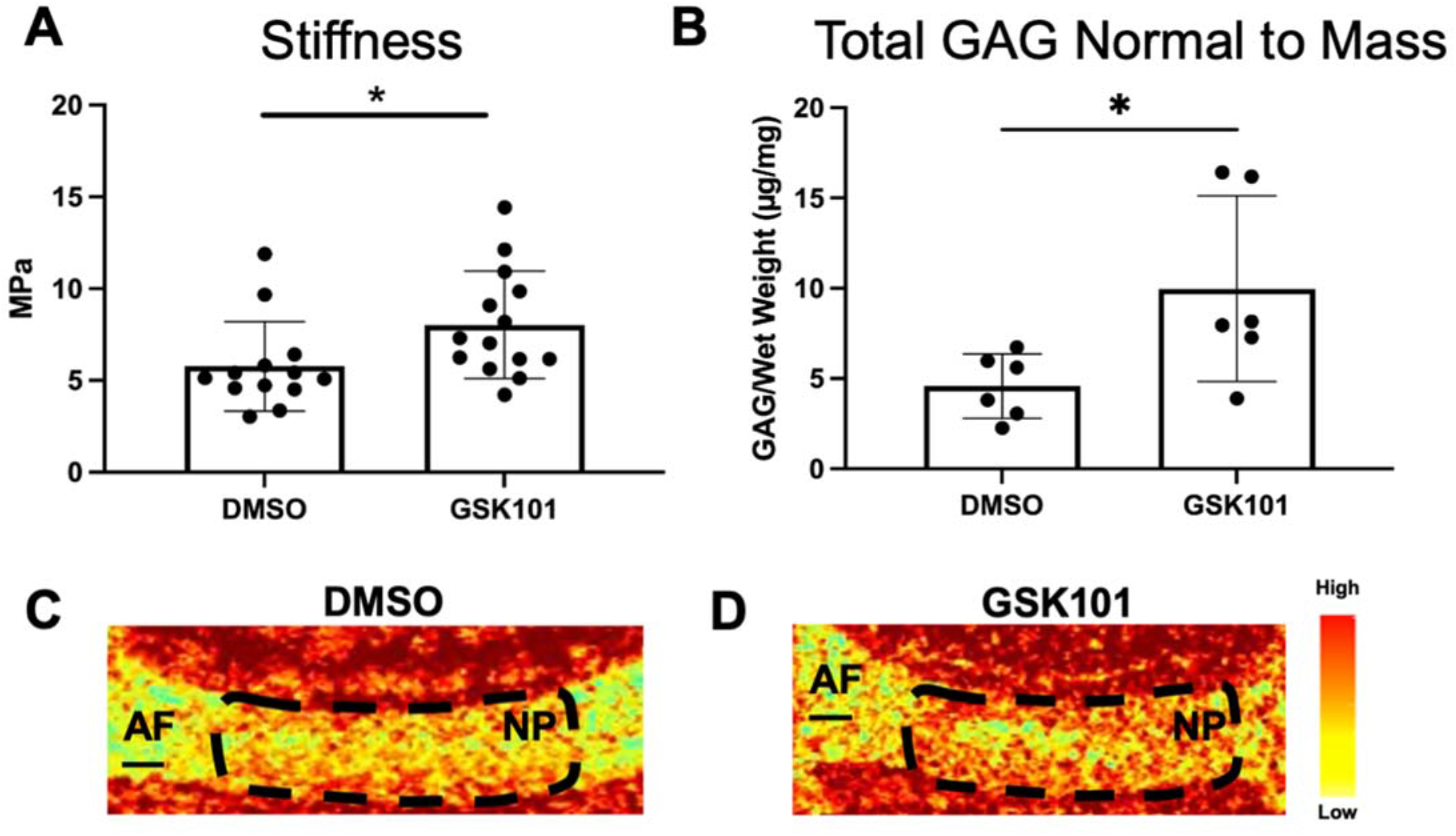
Administering the TRPV4 agonist GSK101 for 7 consecutive days for 3 hours each day in the IVDs resulted in (A) an increase in average stiffness (p<0.05) measured by dynamic compression and (B) an increase in total GAG content in the IVD (p<0.05) measured by the DMMB assay. (C,D) Contrast enhanced microCT (CEμCT) showed that TRPV4-agonized IVDs have high-attenuating nucleus pulposus indicating that there is increased hydration due to the higher GAG composition observed in (B). Data in (A) and (B) were statistically analyzed by ANOVA where * indicates p < 0.05. Scale bars in (C) and (D) are 200μm.

### 3.4 TRPV4 inhibition has no dramatic effect on hypo-osmolarity induced IL-6 production

FSUs were exposed to hypo-osmolar media (200mOsm) for five consecutive days for three hours each day, which significantly increased the production of IL-6 into the culture media at the end of the loading period (Figure 5). In those FSUs where the hypo-osmolarity was co-administered with the TRPV4 inhibitor, GSK205, the IL-6 cytokine concentration in the media was reduced by nearly 50% but not statistically significant (p=0.09).

**Figure 5:**
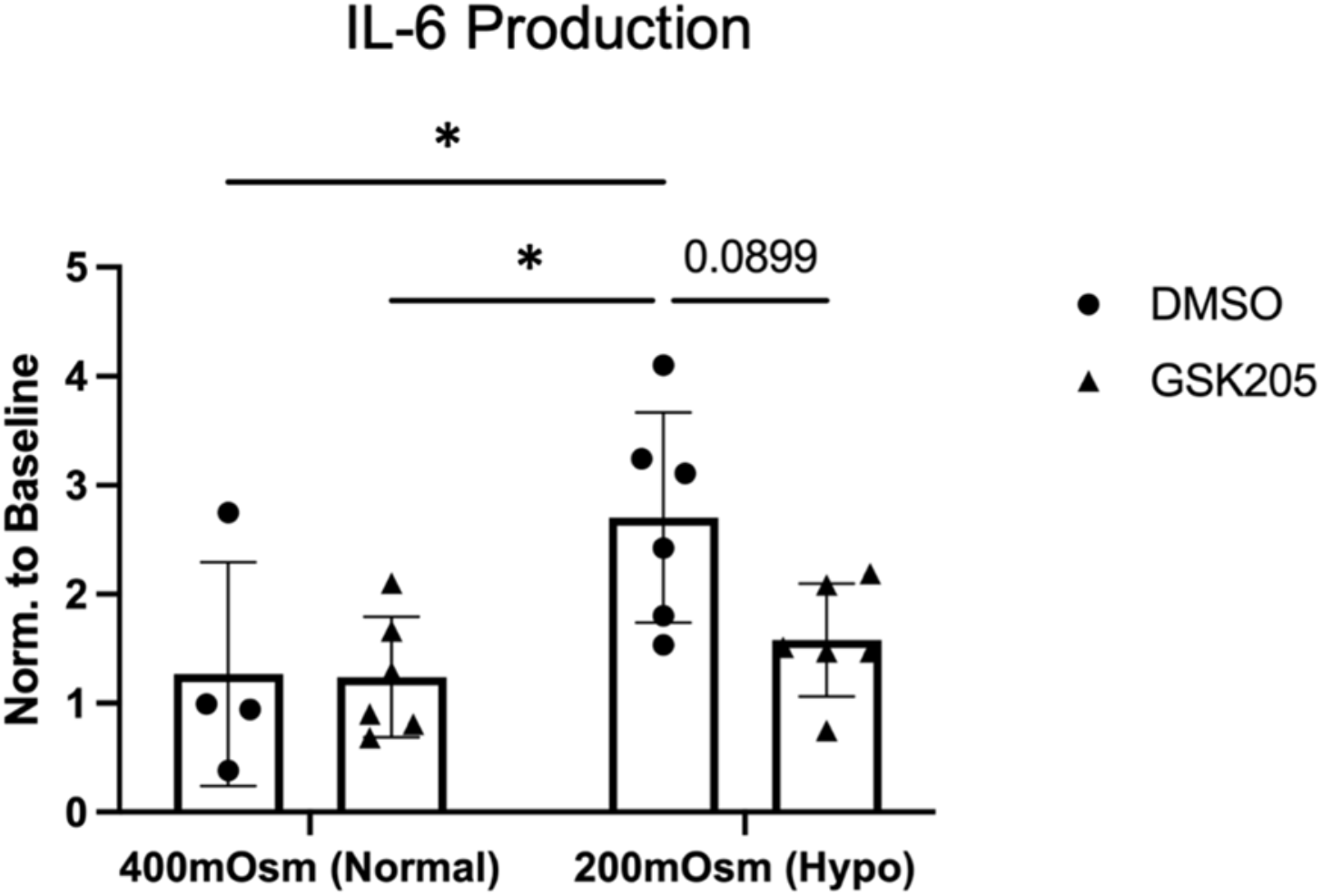
Exposing the IVD to hypo-osmolar pressures during culture significantly increased IL-6 production (p<0.05). The elevated IL-6 production was not significantly resolved with the inhibition of TRPV4 by GSK205 (p = 0.09). Data were statistically analyzed via two-way ANOVA; post-hoc comparisons made by Tukey’s test.

### 3.5 Static loading induced inflammation is partially resolved by TRPV4 inhibition

To evaluate the role of TRPV4 in the IVD load-mediated response, FSUs were observed for 14 days following a 24-hour static loading regimen. The loaded FSUs exhibited significant increase of NF-κB signaling at day 3 of the organ culture compared to the non-loaded control (Figure 6A), through the inhibition of TRPV4 only modestly suppress this NF-κB elevation (Figure 6B). However, by day 7, NF-κB signaling was blunted with TRPV4 inhibition (Figure 6C). Further, concentrations of IL-6 and VEGF-A were both significantly increased at Days 7 and 14 of culture in the loaded FSUs as compared to control (Figure 6C). Notably, IL-6 concentration was significantly lower with TRPV4 inhibition at both Days 7 and 14. The inhibition of TRPV4 had no effect on VEGFA concentration at Day 7, however, it significantly reduced VEGFA at Day 14 compared to the loading condition without GSK205.

**Figure 6:**
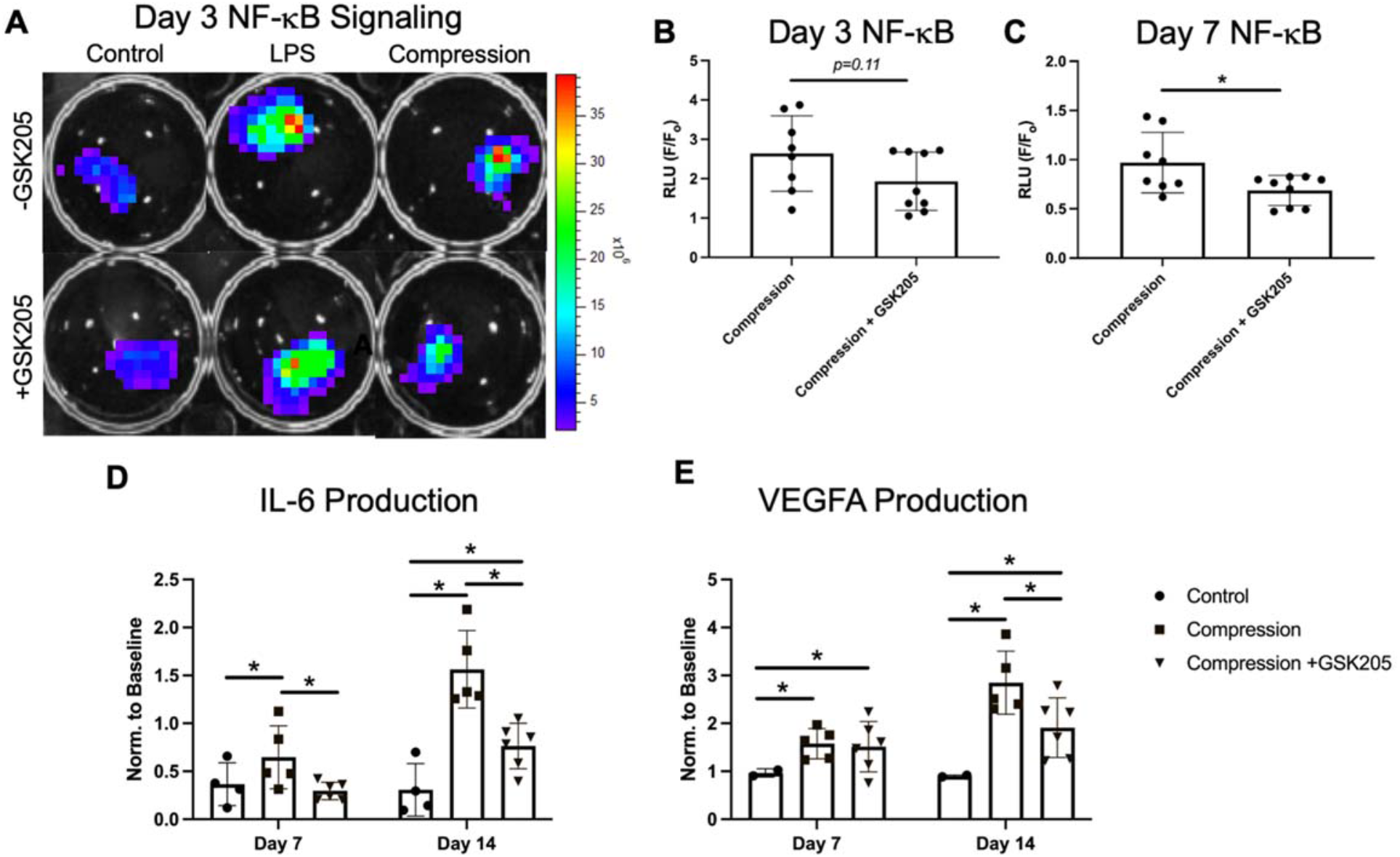
(A) Statically loaded IVDs (“Compression”) exhibited higher NF-κB activity than the unloaded control IVDs. Lipopolysaccharide was used as a positive control to induce the NF-κB luciferase activity. (B,C) NF-κB luciferase activity was reduced with inhibition TRPV4 by GSK205 (Day 3, p=0.11; Day 7, p<0.05). (D, E) The IVDs subjected to compression had increased levels of secreted IL-6 and VEGFA (p<0.05) at both Day 7 and Day 14 of culture. The increase was significantly reduced by TRPV4 inhibition at Day 7 for IL-6 and Day 14 for both IL-6 and VEGFA (p<0.05). Values were normalized to Day 0 media. Data in (B) and (C) were statistically analyzed via unpaired t-test where * indicates p < 0.05. Data in (D) and (E) were statistically analyzed via one-way ANOVA with post hoc Tukey’s HSD where * indicates p < 0.05.

Histological evaluation of compressed IVD revealed a deterioration of the NP-AF boundary, rounded AF cells, and moderate fibrosis in the NP (Figure 7A - F). TRPV4 inhibition by GSK205 in compressed IVDs maintained the distinct boundary between the NP and AF and reduced the number of rounded AF cells (a hall mark of degeneration) (Figure 7BCEF). The degeneration of the IVDs were quantified using a standardized scoring system for mouse IVDs.^40^ IVDs that underwent compression exhibited moderate-to-high degeneration that was significantly higher than control IVDs (Figure 7G). The compressed IVDs with TRPV4 inhibition exhibited mild degeneration. Despite these degenerative characteristics in comparison to the control IVDs, the degeneration score was significantly lower than the compressed IVDs with intact TRPV4 function. Using the histologic scoring system, the individual components of the IVD were assessed (Figure 7H). The IVDs with TRPV4-inhibition had lower degenerative scores for the end plates, NP-AF boundary, and the AF. However, there was little change in the NP degenerative score between the two compressed groups.

**Figure 7:**
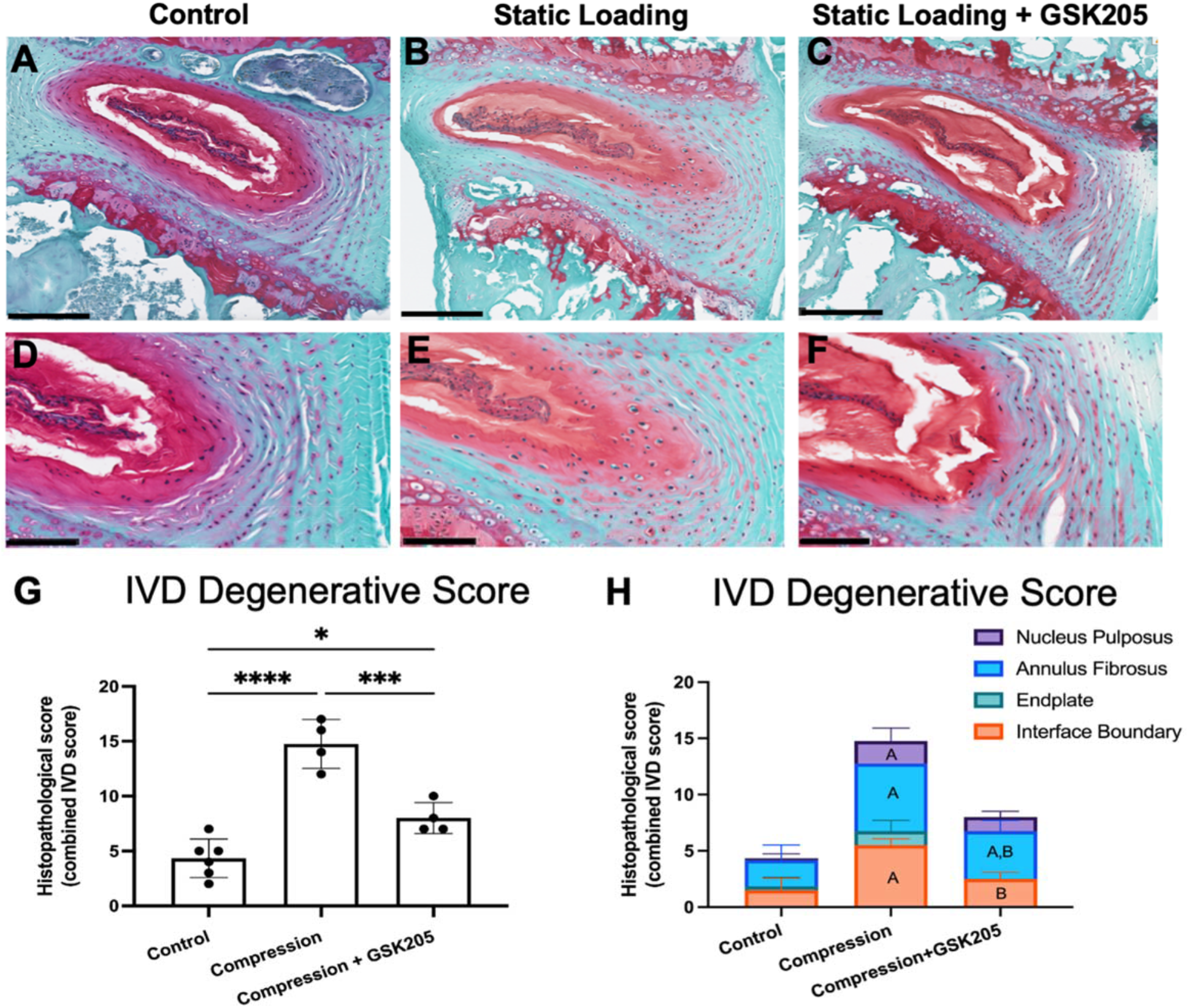
(A - C) The Safranin-O stained histological sections revealed degenerative changes in the compressed IVDs (14-days of loading) that were absent in the non-loaded IVDs. Compressed IVDs with TRPV4 inhibition were partially protected from these degenerative changes. Scale bars are 200μm. (D-F) Qualitatively, the most apparent changes in the loaded IVDs were cell morphology of inner AF cells as well as the boundary between the AF and NP. (G) The degenerative changes in the loaded IVDs were significantly higher than both the controls and the TRPV4 inhibition groups. (H) The protection against degeneration in the TRPV4-inhibited IVDs was primarily in the endplate as well as the boundary interfaces of the IVD (A indicates difference with control group; B indicates differences with compression group). Data in (G,H) were statistically analyzed via ANOVAs respectively with post hoc Tukey’s HSD where * indicates p < 0.05. Scale bars are 100μm.

### 3.6 TRPV4 inhibition does not protect against loading induced collagen disorganization

Quantitative polarized light imaging (QPLI) was used to evaluate the organization of collagen fibers in the AF through the measurements of degree of linear polarization (DoLP) and the angle of polarization (AoP) (Supplemental Figures 3 and 4). DoLP measures the strength of the alignment of collagen fibers and is related to the local retardance and thus structural anisotropy of the tissue: a DoLP of 1 indicates a uniformly aligned tissue whereas a DoLP of 0 indicates isotropic collagen organization. AoP values provide a measurement of the orientation of collagen fibers. Healthy AF can be characterized by a highly organized, lamellae structure of collagen fibers. Inspection of the AoP shows alternating layers of collagen fiber directionality, indicating the primary collagen fiber directions of the alternating lamellae. There were no observed qualitative changes in the AoP or fiber orientation between groups as each showed distinct layers of alternating alignment of approximately equal thickness and spacing (Figure 8A). The compressed IVDs had significantly (p<0.05) decreased average DoLP (AVG DoLP) compared to uncompressed controls, indicating a less strongly aligned tissue (Figure 8B). Compressed IVDs with TRPV4 inhibited did not have significantly decreased average DoLP. In addition, compression decreased the standard deviation of the DoLP (STD DoLP) regardless of TRPV4 inhibition (Figure 8C), indicating decreased uniformity of the collagen fiber alignment in the IVD. Similar levels of STD DoLP for both compression groups indicate similar lamellar tissue structure with or without TRPV4 inhibition. However, inhibiting TRPV4 in compressed IVDs did not rescue collagen disorganization, suggesting that the effects on collagen architecture may be a direct mechanical consequence of loading rather than due to TRPV4-mediated responses.

**Figure 8:**
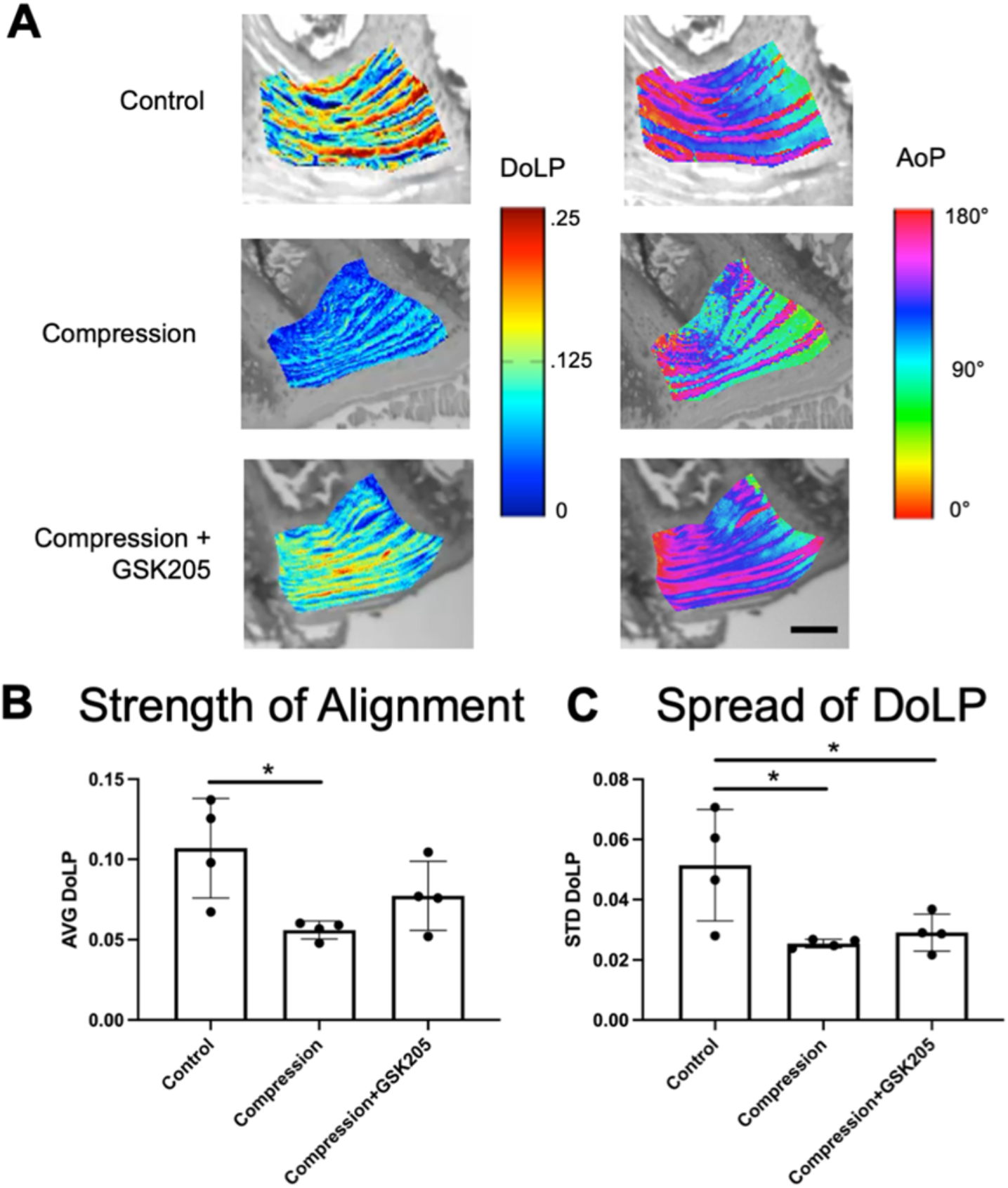
(A) Representative color maps from Quantitative Polarized Light Imaging (QPLI)showing the degree of linear polarization (DoLP) and the angle of polarization (AoP) for control, compression, and compression + GSK205 groups. The DoLP represent the strength of collagen fiber alignment, and the AoP is indicative of fiber orientation. Scale bar is 200μm. (B) Average (AVG) DoLP, showing the average strength of alignment, was decreased within the AF-spanning ROI in the compression group compared to controls. When TRPV4 was inhibited via GSK205, there was no observable difference in AVG DoLP compared to controls. (C) Standard deviation (STD) DoLP, representing the relative spread in DoLP values, was decreased in both experimental groups compared to controls. Data in (B) and (C) were statistically analyzed via one-way ANOVA with post hoc Tukey’s HSD where * indicates p < 0.05.

## Discussion

Our results indicate TRPV4 plays a role in mechanotransduction in response to physiologically relevant loading of the IVD. The effects of TRPV4 on the IVDs depended on the repetition, concentration, and loading history. TRPV4 activation by the small molecule agonist GSK101 in native IVD tissue rapidly initiated Ca^2+^ signaling in AF cells in a dose-dependent manner. NF-κB activity and subsequent IL-6 production also increased following transient activation of TRPV4. The repeated activation of TRPV4 had positive effects in the ECM of the IVD, including improved whole-IVD compressive stiffness and increased glycosaminoglycans in the NP. The prolonged daily repeated exposure to GSKS101 sustained the increase in TRPV4-related outcomes.^27^ Additionally, exposing the IVD to repetitive hypo-osmolar conditions resulted in IL-6 production similar to a single bout of TRPV4 activation. Suppression of TRPV4 concomitant with the hypo-osmolar exposure of the IVDs reduced the IL-6 synthesis.

The sustained physiologic compression of the whole IVD recapitulated some aspects of TRPV4 activation, including NF-κB signaling and IL-6 production. However, sustained compression also created hallmarks of degeneration and microstructural disorganization.^40^ Applying a TRPV4 antagonist during sustained compression partially alleviated the degeneration and the matrix disorganization. Production of cytokines such as VEGFA likely exacerbated any mechanical damage that occurred within the IVD. There were also notable regional variations in degeneration. In particular, TRPV4 inhibition appeared more effective in regions of higher endogenous expression of TRPV4,^34^ with the greatest protective effect in the inner AF. Yet TRPV4 inhibition only conferred partial protection from the degeneration in the outer AF and the interface boundaries. Qualitatively, polarized light imaging revealed greater disorganization in the AF in compressed IVDs.^41^ TRPV4 inhibition did not improve collagen alignment of the compressed IVDs. Along with histopathology scoring, the results suggest that the timely inhibition of TRPV4 can dampen the inflammatory effects on degeneration but not load-mediated mechanical damage. It is worth noting that the inhibition of TRPV4 for seven days depressed NF-κB signaling beyond the baseline, and since NF-kB has critical roles in cell survival, the long-term suppression of NF-kB via inhibiting TRPV4 should be applied cautiously

Similar to articular chondrocytes, activation of TRPV4 in the IVD promoted robust NF-κB signaling that subsided in 24 hours,^44,45^ which is a downstream target of CaMKII and calcium signaling following TRPV4 activation.^46–49^ Though the precise role ofNF-κB is not known here, NF-κB is canonically required to initiate a regenerative response during tissue repair.^50–52^ Consistent with these observations, we observed a spike in NF-κB activity with TRPV4 activation and sustained mechanical loading, followed by improvements in mechanical function and greater GAG density in the NP. Similar to NF-κB, IL-6 can invoke reparative responses in multiple tissues,^53–55^ and it has pleiotropic effects on a myriad of cell types including immune cells and neurons. Similar to our observations, IL-6 has been reported to promote GAG synthesis in fibroblasts.^56^ Conversely, the persistent elevation ofIL-6 and other inflammatory factors is linked to chronic painful intervertebral disc diseases and promote ECM catabolism.^57–59^ Our data here suggest that the TRPV4-dependent activation of IL-6 results in a net increase in GAGs between these known anabolic and catabolic functions. Hypo-osmotic conditions in IVD cells has also stimulated IL-6 expression independently of the TRPV4-pathway, indicating multiple compensatory mechanisms can activate IL-6.^60^

TRPV4 is a critical osmotic and mechanical regulator in multiple organs systems and cell types,^24–26^ and we demonstrate here that it robustly responds to mechanical loading in the intact IVD. There are also a number of known mechanoreceptors, including the TRP family and PIEZO ion channels that regulate of Ca^2+^ flux, that could participate in IVD’s adaptation to sustained loading.^61,62^ In particular, the small molecule inhibitor GSK205 also cross-reacts with TRPA1, inhibiting the ion channel, and other ion channels may have contributed to the observed effects reported in this study.^63^ However, it is worth noting that TRPA1 is primarily expressed in sensory neurons and not in healthy mature IVDs, and therefore is unlikely to be present in the IVDs or influence our *ex vivo* cultures.^64,65^ Moreover, our organ approach here allowed us to selectively isolate and control the IVD cells in their native ECM, ensuring the coupling of our observations between sustained mechanical loading, TRPV4 inhibition, and cellular responses on the IVD tissue. Our results here suggest that TRPV4 is a crucial target for the mechano-responsiveness of the IVD and that modulation of TRPV4 may provide a potential therapeutic target against deleterious loading.

## Acknowledgments

The authors would like to thank Sara Oswald for critical comments and technical editing of the manuscript.

## Author Contributions

Easson, Tang, Lake and Guilak conceived and designed the research; Easson, Savadipour, Anandarajah and Iannucci performed the research and acquired the data; Easson and Iannucci analyzed and interpreted the data. All authors were involved in drafting and revising the manuscript.

## Data Availability Statement

The data that support the findings of this study are available in the methods and/or supplementary material of this article.

## References

1. Balagué F, Mannion AF, Pellisé F, Cedraschi C. Non-specific low back pain. The Lancet. 2012;379(9814):482–491. doi:10.1016/S0140-6736(11)60610-7

2. Maher C, Underwood M, Buchbinder R. Non-specific low back pain. The Lancet. 2017;389(10070):736–747. doi:10.1016/S0140-6736(16)30970-9

3. Jensen M, Brant-Zawadzki M, Obuchowski N, Modic M, Malkasian D, Ross J. Magnetic resonance imaging ofthe lumbar spine in people without back pain. NEJM. 1994;331(2):69–73.

4. Stokes IAF, latridis JC. Mechanical Conditions That Accelerate Intervertebral Disc Degeneration: Overload Versus Immobilization: Spine. 2004;29(23):2724–2732. doi:10.1097/01.brs.0000146049.52152.da

5. latridis JC, MacLean JJ, Roughley PJ, Alini M. Effects of Mechanical Loading on Intervertebral Disc Metabolism In Vivo. Journal of Bone and Joint Surgery. 2006;88(suppl_2):41–46. doi:10.2106/JBJS.E.01407

6. Luoma K, Riihimäki H, Luukkonen R, Raininko R, Viikari-Juntura E, Lamminen A. Low Back Pain in Relation to Lumbar Disc Degeneration: Spine. 2000;25(4):487–492. doi:10.1097/00007632-200002150-00016

7. Fearing BV, Hernandez PA, Setton LA, Chahine NO. Mechanotransduction and cell biomechanics ofthe intervertebral disc. Published online 2018:38.

8. Risbud MV, Shapiro IM. Role of Cytokines in Intervertebral Disc Degeneration: Pain and Disc-content. Published online 2015:24.

9. Chan SCW, Ferguson SJ, Gantenbein-Ritter B. The effects of dynamic loading on the intervertebral disc. European Spine Journal. 2011;20(11):1796–1812. doi:10.1007/s00586-011-1827-1

10. Neidlinger-Wilke C, Galbusera F, Pratsinis H, et al. Mechanical loading of the intervertebral disc: from the macroscopic to the cellular level. European Spine Journal. 2014;23(S3):333–343. doi:10.1007/s00586-013-2855-9

11. Korecki CL, MacLean JJ, latridis JC. Dynamic Compression Effects on Intervertebral Disc Mechanics and Biology: Spine. 2008;33(13):1403–1409. doi:10.1097/BRS.0b013e318175cae7

12. Gawri R, Rosenzweig DH, Krock E, et al. High mechanical strain of primary intervertebral disc cells promotes secretion of inflammatory factors associated with disc degeneration and pain. Arthritis Research & Therapy. 2014;16(1):R21. doi:10.1186/ar4449

13. Scheller J, Chalaris A, Schmidt-Arras D, Rose-John S. The pro-and anti-inflammatory properties of the cytokine interleukin-6. Biochimica et Biophysica Acta (BBA) - Molecular Cell Research. 2011;1813(5):878–888. doi:10.1016/j.bbamcr.2011.01.034

14. McKenzie JA, Bixby EC, Silva MJ. Differential Gene Expression from Microarray Analysis Distinguishes Woven and Lamellar Bone Formation in the Rat Ulna following Mechanical Loading. Beier F, ed. PLoS ONE. 2011;6(12):e29328. doi:10.1371/journal.pone.0029328

15. Johnson BZ, Stevenson AW, Prêle CM, Fear MW, Wood FM. The Role of IL-6 in Skin Fibrosis and Cutaneous Wound Healing. Biomedicines. 2020;8(5):101. doi:10.3390/biomedicines8050101

16. Wiet MG, Piscioneri A, Khan SN, Ballinger MN, Hoyland JA, Purmessur D. Mast Cell-Intervertebral disc cell interactions regulate inflammation, catabolism and angiogenesis in Discogenic Back Pain. Sci Rep. 2017;7(1):12492. doi:10.1038/s41598-017-12666-z

17. Tian Y, Yuan W, Fujita N, et al. Inflammatory Cytokines Associated with Degenerative Disc Disease Control Aggrecanase-1 (ADAMTS-4) Expression in Nucleus Pulposus Cells through MAPK and NF-κB. The American Journal of Pathology. 2013;182(6):2310–2321. doi:10.1016/j.ajpath.2013.02.037

18. Ebbinghaus M, Uhlig B, Richter F, et al. The role of interleukin-1β in arthritic pain: Main involvement in thermal, but not mechanical, hyperalgesia in rat antigen-induced arthritis. Arthritis & Rheumatism. 2012;64(12):3897–3907. doi:10.1002/art.34675

19. Kim CF, Moalem-Taylor G. lnterleukin-17 Contributes to Neuroinflammation and Neuropathic Pain Following Peripheral Nerve Injury in Mice. The Journal of Pain. 2011;12(3):370–383. doi:10.1016/j.jpain.2010.08.003

20. Martinac B. Mechanosensitive ion channels: molecules of mechanotransduction. Journal of Cell Science. 2004;117(12):2449–2460. doi:10.1242/jcs.01232

21. Venkatachalam K, Montell C. TRP Channels. Annual Review of Biochemistry. 2007;76(1):387–417. doi:10.1146/annurev.biochem.75.103004.142819

22. Everaerts W, Nilius B, Owsianik G. The vanilloid transient receptor potential channel TRPV4: From structure to disease. Progress in Biophysics and Molecular Biology. 2010;103(1):2–17. doi:10.1016/j.pbiomolbio.2009.10.002

23. White JPM, Cibelli M, Urban L, Nilius B, McGeown JG, Nagy I. TRPV4: Molecular Conductor of a Diverse Orchestra. Physiol Rev. 2016;96:63.

24. Poole K. The Diverse Physiological Functions of Mechanically Activated Ion Channels in Mammals. Annu Rev Physiol. 2022;84(1):307–329. doi:10.1146/annurev-physiol-060721-100935

25. Thorneloe KS, Cheung M, Bao W, et al. An Orally Active TRPV4 Channel Blocker Prevents and Resolves Pulmonary Edema Induced by Heart Failure. Sci Transl Med. 2012;4(159). doi:10.1126/scitranslmed.3004276

26. Wu L, Gao X, Brown RC, Heller S, O’Neil RG. Dual role of the TRPV4 channel as a sensor of flow and osmolality in renal epithelial cells. American Journal of Physiology-Renal Physiology. 2007;293(5):F1699–F1713. doi:10.1152/ajprenal.00462.2006

27. O’Conor CJ, Leddy HA, Benefield HC, Liedtke WB, Guilak F. TRPV4-mediated mechanotransduction regulates the metabolic response of chondrocytes to dynamic loading. Proceedings of the National Academy of Sciences. 2014;111(4):1316–1321. doi:10.1073/pnas.1319569111

28. O’Conor CJ, Ramalingam S, Zelenski NA, et al. Cartilage-Specific Knockout of the Mechanosensory Ion Channel TRPV4 Decreases Age-Related Osteoarthritis. Scientific Reports. 2016;6(1). doi:10.1038/srep29053

29. Phan MN, Leddy HA, Votta BJ, et al. Functional characterization of TRPV4 as an osmotically sensitive ion channel in porcine articular chondrocytes. Arthritis & Rheumatism. 2009;60(10):3028–3037. doi:10.1002/art.24799

30. Pritchard S, Erickson GR, Guilak F. Hyperosmotically Induced Volume Change and Calcium Signaling in Intervertebral Disk Cells: The Role of the Actin Cytoskeleton. Biophysical Journal. 2002;83(5):2502–2510. doi:10.1016/S0006-3495(02)75261-2

31. Pritchard S, Guilak F. The Role of F-Actin in Hypo-Osmotically Induced Cell Volume Change and Calcium Signaling in Anulus Fibrosus Cells. Annals of Biomedical Engineering. 2004;32(1):103–111. doi:10.1023/B:ABME.0000007795.69001.35

32. Walter BA, Purmessur D, Moon A, et al. Reduced tissue osmolarity increases TRPV4 expression and pro-inflammary cytokines in intervertebral disc cells. Published online 2016:21.

33. Cambria E, Arlt MJE, Wandel S, et al. TRPV4 Inhibition and CRISPR-Cas9 Knockout Reduce Inflammation Induced by Hyperphysiological Stretching in Human Annulus Fibrosus Cells. Cells. 2020;9(7):1736. doi:10.3390/cells9071736

34. Kim M, Ramachandran R, Seguin C. Spatiotemporal and functional characterisation of transient receptor potential vanilloid 4 (TRPV4) in the murine intervertebral disc. eCM. 2021;41:194–203. doi:10.22203/eCM.v041a14

35. Liu JW, Lin KH, Weber C, et al. An In Vitro Organ Culture Model ofthe Murine Intervertebral Disc. JoVE. 2017;(122):55437. doi:10.3791/55437

36. Orrenius S, Zhivotovsky B, Nicotera P. Regulation of cell death: the calcium–apoptosis link. Nat Rev Mol Cell Biol. 2003;4(7):552–565. doi:10.1038/nrm1150

37. Lin KH, Wu Q, Leib DJ, Tang SY. A novel technique for the contrast-enhanced microCT imaging of murine intervertebral discs. Journal of the Mechanical Behavior of Biomedical Materials. 2016;63:66–74. doi:10.1016/j.jmbbm.2016.06.003

38. Liu JW, Abraham AC, Y. Tang S. The high-throughput phenotyping of the viscoelastic behavior of whole mouse intervertebral discs using a novel method of dynamic mechanical testing. Journal of Biomechanics. 2015;48(10):2189–2194. doi:10.1016/j.jbiomech.2015.04.040

39. Wilke HJ, Neef P, Caimi M, Hoogland T, Claes L. New In Vivo Measurements of Pressures in the Intervertebral Disc in Daily Life. Spine. 1995;24(8):755–762.

40. Melgoza IP, Chenna SS, Tessier S, et al. Development of a standardized histopathology scoring system using machine learning algorithms for intervertebral disc degeneration in the mouse model—An ORS spine section initiative. JOR Spine. 2021;4(2). doi:10.1002/jsp2.1164

41. York T, Kahan L, Lake SP, Gruev V. Real-time high-resolution measurement of collagen alignment in dynamically loaded soft tissue. J Biomed Opt. 2014;19(06):1. doi:10.1117/1.JBO.19.6.066011

42. lannucci L, Riak MB, Lake SP. The effect of extracellular matrix properties on polarized light-based analysis of collagen fiber alignment in soft tissues. In: Ramella-Roman JC, Ma H, Vitkin IA, Elson DS, Novikova T, eds. Polarized Light and Optical Angular Momentum for Biomedical Diagnostics 2022. SPIE; 2022:4. doi:10.1117/12.2614438

43. Lin KH, Tang SY. The Quantitative Structural and Compositional Analyses of Degenerating Intervertebral Discs Using Magnetic Resonance Imaging and Contrast-Enhanced Micro-Computed Tomography. Ann Biomed Eng. 2017;45(11):2626–2634. doi:10.1007/s10439-017-1891-8

44. Nims R, Pferdehirt L, Ho N, et al. A synthetic mechanogenetic gene circuit for autonomous drug delivery in engineered tissues. SCIENCE ADVANCES. Published online 2021:14.

45. Nims RJ, Pferdehirt L, Guilak F. Mechanogenetics: harnessing mechanobiology for cellular engineering. Current Opinion in Biotechnology. 2022;73:374–379. doi:10.1016/j.copbio.2021.09.011

46. Kashiwase K, Higuchi Y, Hirotani S, et al. CaMKII activates ASK1 and NF-κB to induce cardiomyocyte hypertrophy. Biochemical and Biophysical Research Communications. 2015;327(1):136–142. doi:10.1016/j.bbrc.2004.12.002

47. Williams KM, Leser JM, Gould NR, et al. TRPV4 calcium influx controls sclerostin protein loss independent of purinergic calcium oscillations. Bone. 2020;136:115356. doi:10.1016/j.bone.2020.115356

48. Lyons JS, Joca HC, Law RA, et al. Microtubules tune mechanotransduction through NOX2 and TRPV4 to decrease sclerostin abundance in osteocytes. Sci Signal. 2017;10(506):eaan5748. doi:10.1126/scisignal.aan5748

49. Martin TP, McCluskey C, Cunningham MR, Beattie J, Paul A, Currie S. CaMKIIδ interacts directly with IKKβ and modulates NF-κB signalling in adult cardiac fibroblasts. Cellular Signalling. 2018;51:166–175. doi:10.1016/j.cellsig.2018.07.008

50. Karra R, Knecht AK, Kikuchi K, Poss KD. Myocardial NF-κB activation is essential for zebrafish heart regeneration. Proc Natl Acad Sci USA. 2015;112(43):13255–13260. doi:10.1073/pnas.1511209112

51. Lin T, Pajarinen J, Nabeshima A, et al. Establishment of NF-κB sensing and interleukin-4 secreting mesenchymal stromal cells as an “on-demand” drug delivery system to modulate inflammation. Cytotherapy. 2017;19(9):10251–1034. doi:10.1016/j.jcyt.2017.06.008

52. Xiong Y, Li W, Shang C, et al. Brg1 Governs a Positive Feedback Circuit in the Hair Follicle for Tissue Regeneration and Repair. Developmental Cell. 2013;25(2):169–181. doi:10.1016/j.devcel.2013.03.015

53. Kang S, Tanaka T, Narazaki M, Kishimoto T. Targeting lnterleukin-6 Signaling in Clinic. Immunity. 2019;50(4):1007–1023. doi:10.1016/j.immuni.2019.03.026

54. Muñoz-Cánoves P, Scheele C, Pedersen BK, Serrano AL. Interleukin-6 myokine signaling in skeletal muscle: a double-edged sword? FEBSJ. 2013;280(17):4131–4148. doi:10.1111/febs.12338

55. Gadient RA, Otten UH. Interleukin-6 (IL-6)—A molecule with both beneficial and destructive potentials. Progress in Neurobiology. 1997;52(5):379–390. doi:10.1016/S0301-0082(97)00021-X

56. Duncan M, Berman B. Stimulation of collagen and glycoaminoglycan production in cultured human adult dermanl fibroblasts by recombinant human interleukin 6. JID. 1991;97(4):686–692.

57. Heffner KL, France CR, Trost Z, Mei Ng H, Pigeon WR. Chronic Low Back Pain, Sleep Disturbance, and Interleukin-6. The Clinical Journal of Pain. 2011;27(1):35–41. doi:10.1097/AJP.0b013e3181eef761

58. Weber KT, Alipui DO, Sison CP, et al. Serum levels ofthe proinflammatory cytokine interleukin-6 vary based on diagnoses in individuals with lumbar intervertebral disc diseases. Arthritis Res Ther. 2016;18(1):3. doi:10.1186/s13075-015-0887-8

59. Rendina D, De Filippo G, Postiglione L, et al. Interleukin-6 trans-signaling and pathological low back pain in patients with Paget disease of bone. Pain. 2018;159(8):1664–1673. doi:10.1097/j.pain.0000000000001260

60. Sadowska A, Altinay B, Hitzl W, Ferguson SJ, Wuertz-Kozak K. Hypo-Osmotic Loading Induces Expression of IL-6 in Nucleus Pulposus Cells of the Intervertebral Disc Independent of TRPV4 and TRPM7. Front Pharmacol. 2020;11:952. doi:10.3389/fphar.2020.00952

61. Lee W, Leddy HA, Chen Y, et al. Synergy between Piezo1 and Piezo2 channels confers high-strain mechanosensitivity to articular cartilage. Proc Natl Acad Sci USA. 2014;111(47). doi:10.1073/pnas.1414298111

62. Lee W, Nims RJ, Savadipour A, et al. Inflammatory signaling sensitizes Piezo1 mechanotransduction in articular chondrocytes as a pathogenic feed-forward mechanism in osteoarthritis. Proc Natl Acad Sci USA. 2021;118(13):e2001611118. doi:10.1073/pnas.2001611118

63. Kanju P, Chen Y, Lee W, et al. Small molecule dual-inhibitors of TRPV4 and TRPA1 for attenuation of inflammation and pain. Scientific Reports. 2016;6(1). doi:10.1038/srep26894

64. Wang XL, Cui LW, Liu Z, et al. Effects of TRPA1 activation and inhibition on TRPA1 and CGRP expression in dorsal root ganglion neurons. Neural Regen Res. 2019;14(1):140. doi:10.4103/1673-5374.243719

65. Kameda T, Zvick J, Vuk M, et al. Expression and Activity of TRPAl and TRPV1 in the Intervertebral Disc: Association with Inflammation and Matrix Remodeling. UMS. 2019;20(7):1767. doi:10.3390/ijms20071767

