## Supplemental Data for "Modulation of TRPV4 Protects against Degeneration Induced by Sustained Loading and Promotes Matrix Synthesis in the Intervertebral Disc"

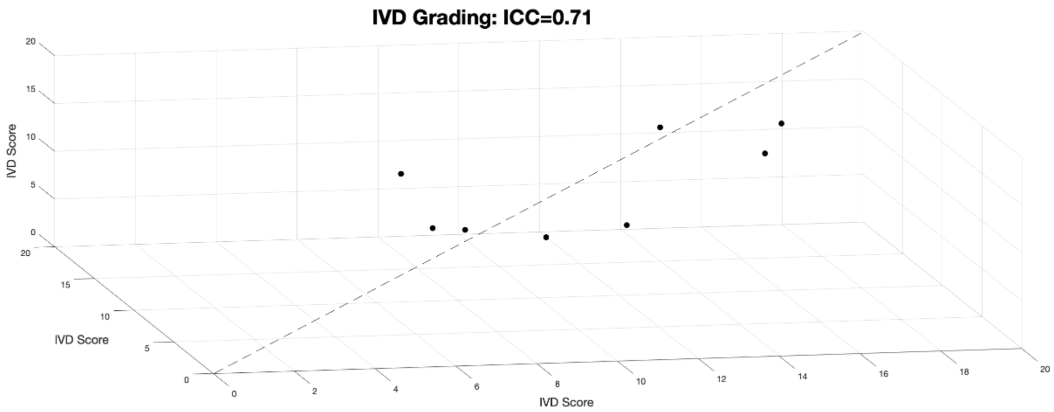


**Supplemental Figure S1:** IVDs were blindly graded using a standardized histological scoring system[40]. A set of IVDs, ranging from healthy to severely degenerated, were graded by a single grader on three non-consecutive days and intraclass correlation (ICC) was measured to be 0.71.

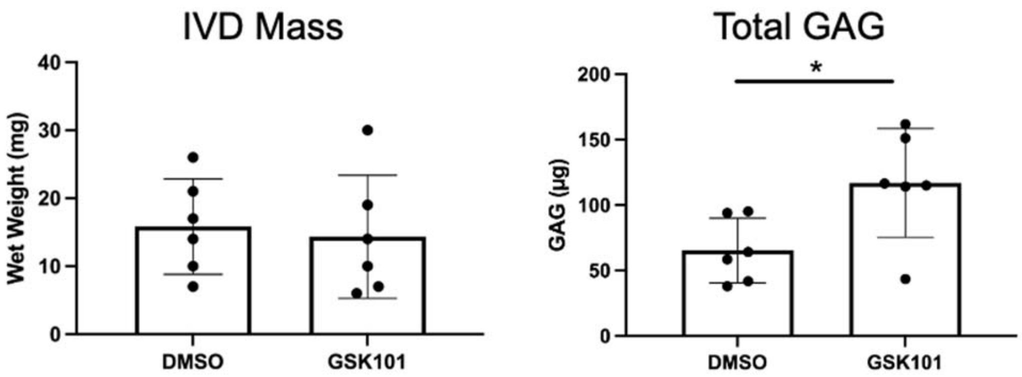


**Supplemental Figure S2:** (A) IVDs were massed following culture and prior to DMMB assay. No differences in wet weights were observed. (B) GAG content was normalized to IVD wet weight. The increased GAG content was significantly higher in the TRPV4-activated IVDs ($p<0.05$). Data were statistically analyzed by student's t-test.

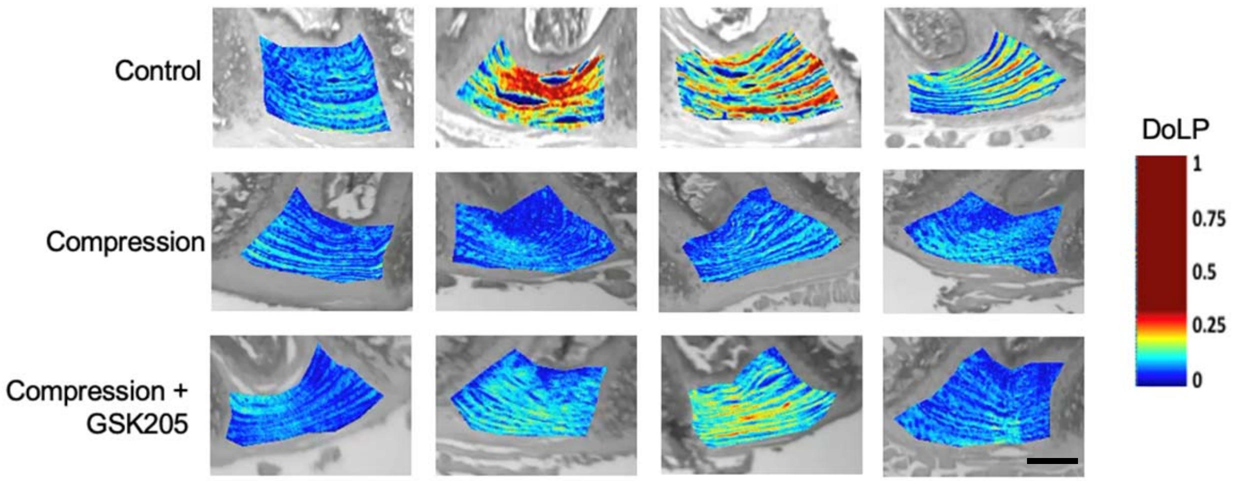


**Supplemental Figure S3:** Degree of linear polarization (DoLP) color maps for all samples analyzed with quantitative polarized light microscopy (QPLI). Not all compressed samples with TRPV4 inhibition (GSK205) were protected against fiber disorganization. Scale bar is 200µm.

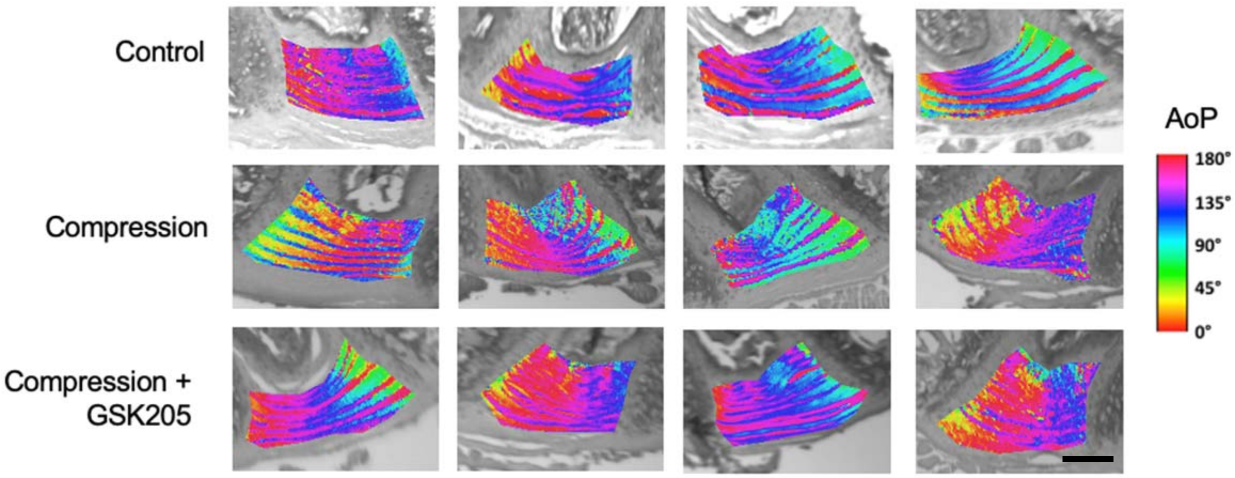


**Figure S4:** Angle of polarization (AoP) color maps for all samples analyzed with quantitative polarized light microscopy (QPLI). Due to variability in fiber angle, AoP was not used to quantify IVD response to static loading. Scale bar is 200µm.
